# MassID provides near complete annotation of metabolomics data with identification probabilities

**DOI:** 10.64898/2026.02.11.704864

**Authors:** Ethan Stancliffe, Monil Gandhi, Douglas V. Guzior, Ashima Mehta, Sandeep Acharya, Adam Richardson, Kevin Cho, Tom Cohen, Gary J. Patti

**Affiliations:** Panome Bio, Saint Louis, MO; Washington University, Saint Louis, MO

## Abstract

Liquid chromatography coupled to mass spectrometry (LC/MS) is a powerful tool in metabolomics research, generating tens-of-thousands of signals from a single biological sample. However, current software solutions for unbiased assessment of metabolomics data analysis are limited by complex sources of noise and non-quantitative metabolite identifications that make results difficult to interpret. Here, we present MassID, a cloud-based untargeted metabolomics pipeline that aims to overcome the innate challenges of unbiased metabolite analysis and perform end-to-end data processing, transforming raw spectra to normalized and identified metabolite profiles. MassID incorporates a suite of software functionalities, including deep learning-based peak detection and comprehensive noise filtering. In addition, with MassID we introduce a novel software module: DecoID2 that enables probabilistic metabolite identification for false discovery rate (FDR)-controlled metabolomics. When applied to a human plasma dataset, MassID results in near-complete signal annotation, identification of >4,000 metabolites (including >1,200 compounds at an FDR <5%) across four complementary LC/MS runs, and enables integrated downstream analyses to understand biochemical dysregulation at both the molecular and pathway level. When compared to the Metabolomics Standards Initiative (MSI) confidence levels, identification probability generally correlated with MSI levels. However, only 356/418 of MSI Level 1 compounds were identified with <5% FDR and the remaining 884 FDR < 5% compounds were identified from MSI L2-L3 compounds, highlighting the enhanced specificity and discovery potential achieved by MassID.

## Introduction

Metabolomics is the study and measurement of metabolites, which are small molecules that play crucial roles in biological systems, including energy production, synthesis of macromolecules, and cellular communication.^1^ Additionally, metabolomics offers the ability to profile lifestyle and exogenous factors that appear as additional small molecules in biospecimens.^2^ The levels of metabolites therefore serve as powerful indicators of cellular processes and disease states, making metabolomics a valuable tool across various fields, including medicine, agriculture, and environmental science. In recent years, metabolomics has been increasingly leveraged in population-scale multi-omic studies, leading to novel understanding in, among others, cancer, aging, metabolic disease, and neurodegeneration.^3–6^

Liquid chromatography coupled to mass spectrometry (LC/MS) is the dominant analytical approach for metabolomics analyses due to its ability to detect a broad range of metabolites with high sensitivity and specificity.^7^ Often, multiple LC/MS methods will be used in combination that operate with different chromatography or ionization polarity to capture the breadth of chemical diversity in the metabolome (e.g., polar metabolites and lipids).^8,9^ Metabolomics can be performed in either an untargeted (looking for all metabolites in the sample) or targeted (looking for specific metabolites) fashion.

In targeted metabolomics, metabolites are searched for using so-called metabolite libraries containing reference data (spectra, retention times, etc.) from authentic standards to identify and quantify the abundance of a compound.^10^ While targeted metabolomics holds value in specific scenarios, its limitations are apparent. The restricted scope of targeted analysis, confined to a predetermined set of metabolites, can lead to overlooked discoveries. Novel metabolites important for human health are frequently being discovered via untargeted analysis, such as 2-hydroxyglutate production in glioma^11^ and Lac-Phe^12^ excretion during exercise. Discovery of novel metabolites is, by definition, impossible with targeted analysis as they will be unknown at the time of analysis. Further, the metabolome encompasses both endogenous metabolites in addition to small molecules and their derivatives taken in from the environment (food, air pollutants, chemical exposures, cosmetics, etc.).^13^ The latest release of the Human Metabolome Database (HMDB)^14^ lists over 200,000 metabolites that have been detected in human biospecimens and the PubChem^15^ database encompasses over 100,000,000 small molecules. In total, this creates a near-infinite chemical space that metabolomics aims to profile, limiting the applicability of targeted analysis for discovery applications.

For unbiased, broad metabolome assessment, software must be employed. Untargeted analysis of metabolomics data typically involves several steps, including peak detection, peak grouping/filtering, metabolite identification, and statistical analysis.^16^ In untargeted metabolomics it is common to detect over 20,000 unique signals from a single LC/MS experiment.^7^ However, recent work has shown that the vast majority of these signals do not provide unique biological information.^17,18,19^

While there are several software tools available for completing different steps of untargeted processing, they often struggle to accurately remove complex sources of chemical noise and to our knowledge, no metabolomics software can provide validated, probabilistic metabolite identifications.^20^ Currently, metabolite identifications are qualitatively scored based on the type of reference data used to make the metabolite identification (MS/MS, RT, external reference data, experimental data, etc.). This scoring scheme, created by the Metabolomics Standards Initiative (MSI), has been instrumental in the progression of metabolomics since its inception in 2007.^21^ However, these scores critically do not provide a quantitative assessment of how likely a metabolite identification is correct. Recently, another approach to metabolite identification scoring was proposed, identification probability, that scores metabolites according to not only matching against reference data but also factoring in alternative matches to compute a probability of each possible metabolite identification.^22^ While a powerful concept, statistical formalism and validation is needed to enable true probabilistic metabolite identifications that enable false discovery rate (FDR)-controlled metabolomics.

To address these challenges, we have developed MassID, a cloud-based untargeted pipeline that performs end-to-end analysis of LC/MS/MS data. MassID incorporates a suite of both novel and existing software functionalities, which leverage deep learning, probabilistic modeling, and other machine learning methods to eliminate noise, quantitatively score metabolite identifications, and minimize technical variability. MassID has been optimized to process LC/MS/MS data from complementary reversed phase and hydrophilic interaction liquid chromatography assays designed to provide broad but efficient coverage of the metabolome. Here we present the MassID software, validation data, and an application to human plasma where more than 4,500 metabolites were identified.

## Results and Discussion

### Metabolomics Workflow

MassID is a software suite optimized for analysis of LC/MS-based metabolomics data generated with both reversed-phase liquid chromatography (RPLC) and hydrophilic interaction liquid chromatography (HILIC) generated in both positive and negative ionization modes, amounting to a total of four LC/MS assays that are applied to each sample. These analytical methods were selected to optimize metabolome coverage while minimizing the number of analytical runs required.^9,23^ The overall data generation and analysis workflow is provided in Figure 1a.

**Figure 1:**
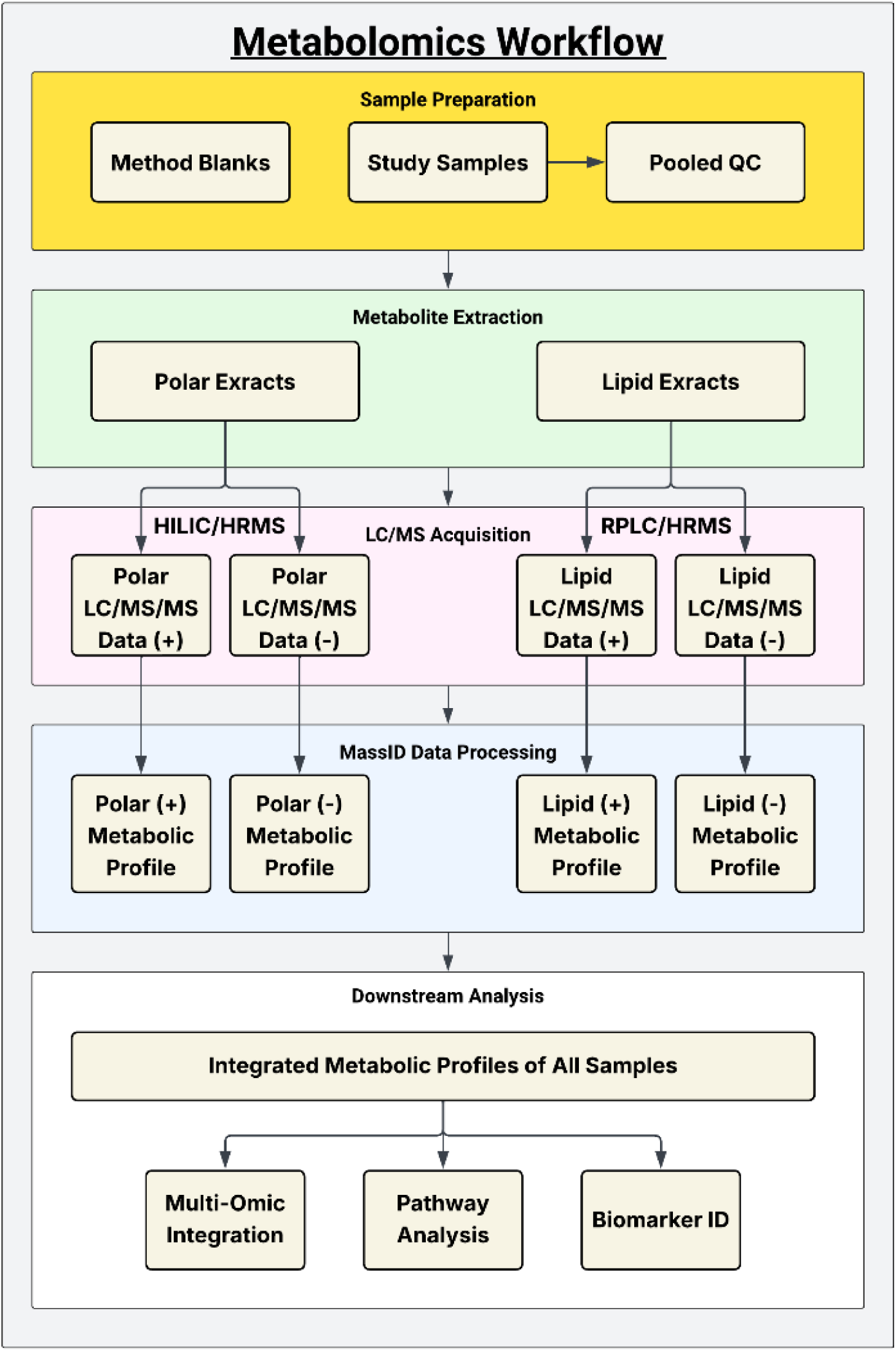
Metabolomics Workflow. LC/MS data is generated through two different chemical extractions and LC methods (RPLC and HILIC). In addition, both positive and negative ionization modes are utilized for analysis. MassID is applied to the resulting data from each of the four assays and the outputs are combined to form an integrated metabolic profile of each sample that can be leveraged for downstream analysis.

First, in addition to the study samples, method blanks and pooled quality control (QC) samples formed by combining a small amount of each study sample are prepared that will be leveraged throughout the workflow. Polar and lipid metabolites are extracted separately for each sample, QC, and blank, leveraging a solid-phase extraction protocol (see Methods). High-resolution mass spectrometry data is acquired in MS1, full scan mode on every sample. The QC sample is injected at the beginning and end of each analysis batch (typically ∼90 samples), as well as in between every 10-12 samples to monitor and correct for batch and run-order effects. Further, the QC sample is utilized for iterative data-dependent MS/MS data acquisition (DDA).^24^ The details of our analytical workflow are provided in the Methods as well as our QC procedures.

After data acquisition, the MassID software is applied to perform an untargeted analysis of the acquired LC/MS/MS data for each of the four assays independently. The output of MassID is processed metabolic profiles containing the relative abundance of both identified and unidentified metabolites across all samples. After MassID processing, the results from all assays are combined to form a single integrated metabolic profile of each study sample that can be fed into downstream analysis workflows for, among others, biomarker discovery, pathway analysis, and multi-omic integration. As some metabolites will be detected across multiple assays, we select the most robust measurement for each metabolite to be included in the final output. This selection is first based on identification confidence, then by technical variability (coefficient of variation across the QC samples).

### MassID Software

To perform the untargeted bioinformatic analysis of the high-resolution LC/MS data generated in the workflow described above, we developed the MassID software suite, which has the functionality to perform elucidation of the study-specific metabolites for a project as well as scalable extraction of the relative abundance of each metabolite across an effectively unlimited number of samples. MassID builds upon many core software modules in the field, while incorporating several novel aspects. The overall MassID processing pipeline is shown in Figure 2 and each module is described in the sections below.

**Figure 2:**
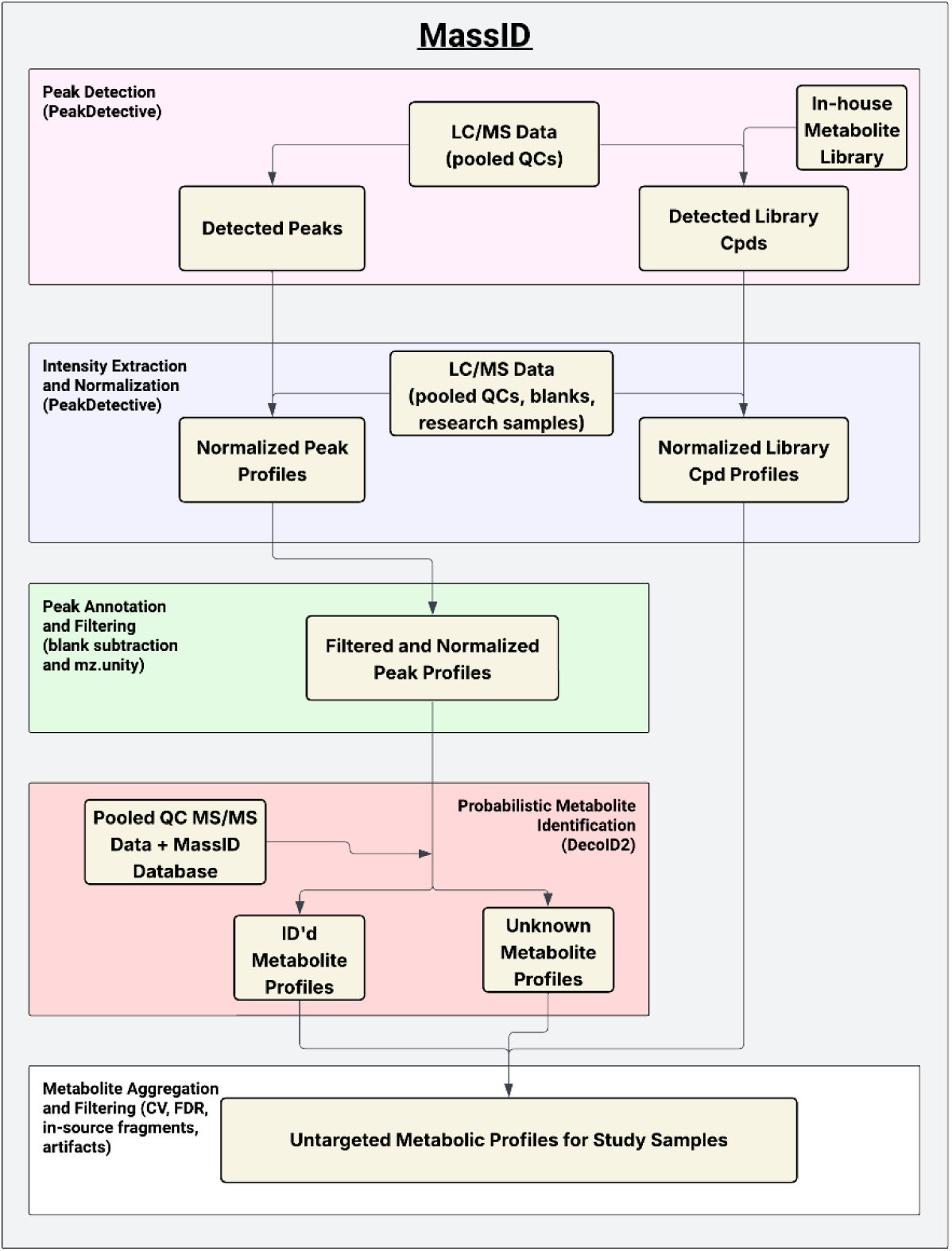
MassID Analysis Pipeline. Peaks are first detected from the LC/MS data from QCs samples. Peak detection is performed in an untargeted fashion (looking for all signals) as well as in a targeted fashion looking for metabolites corresponding to compounds from the metabolite library. Following peak detection, the untargeted peak profiles are extracted from all research samples and method blanks, missing values are imputed, and intensities are batch corrected. Subsequently, features are filtered, and remaining features are subject to metabolite identification through the application of the novel DecoID2 software. Lastly, final filtering of profiled metabolites are filtered based on CV, FDR, and in-source fragments. The filtered metabolic profiles of identified and unknown metabolites are then combined with the detected library compounds to form the metabolic profiles of the study samples, QCs, and blanks.

### Peak Detection

For detecting metabolite signals from the LC/MS data, PeakDetective^25^, a deep learning-based peak detection algorithm, is used to elucidate metabolite signals with greater sensitivity and specificity than conventional approaches. However, PeakDetective is computationally expensive when leveraged for peak detection. Therefore, to reduce the analysis time, we perform peak detection only on the pooled QC samples, which represent a chemical average of all study samples. Previous work has confirmed that pooled QC samples enable detection of the majority of metabolites contained within individual study samples.^26^ In MassID, peak detection is performed twice, once in an untargeted fashion looking for all metabolite signals and once in a targeted fashion, looking for compounds present within our in-house library. While most of the library compounds will already be detected with the untargeted peak detection, targeted analyses can increase sensitivity in some cases.^10^

### Intensity Extraction and Normalization

After both targeted and untargeted peak detection, the metabolite peak areas are extracted from the raw data files of all study samples, QCs, and blanks based on the *m/z* and retention time coordinates of each peak. Retention times are aligned (via dynamic time warping)^27,28^ to the reference QC samples and uniform integration bounds are applied based on the average full-width half maximum peak boundaries.

To reduce technical variation and enable rigorous downstream statistical analysis, post-processing of the metabolite intensities is performed. First, missing values are imputed. Depending on the data processing procedure applied, missing values can occur due to systematic or random causes. For MassID, metabolite intensities are extracted from the data based on defined *m/z* and retention time windows, which creates missing values that are largely due to metabolites falling below the limit of detection of the instrument. As such, we apply a half-minimum imputation method where missing values are imputed per metabolite with half the minimum detected intensity for the compound. This methodology has been previously shown to achieve high accuracy for missing values caused primarily by metabolites falling below the LOD.^29,30^

In addition to missing values, technical variation that can be attributed to batch or run-order effects is removed via a random forest-based methodology we previously reported.^26,31^ In brief, the intensity drift per metabolite in the QC samples (analyzed throughout the study) is calculated and a random forest model is fit to this drift using analysis batch and injection order as predictors. After fitting, the predicted drift per metabolite is estimated for all research samples and subtracted from the metabolite intensities.

Lastly, metabolite intensities for all samples are normalized following a delayed normalization approach^32^ that is commonly applied in bottom-up proteomics to account for sample-wide effects from pipetting or other sample handling errors. This methodology finds a scaling factor applied to all metabolites in a sample to minimize differences in median signal intensity across all samples. After normalization, metabolite intensities are log2 transformed to enable downstream statistical analysis.

For peaks found from untargeted detection, further steps must be applied to annotate, filter, and identify peaks prior to final data output. For metabolites found from targeted detection, after imputation and data normalization, the resulting metabolite intensities are ready for downstream analysis.

### Peak Annotation and Filtration

Once untargeted peak detection has been completed, annotation of these peaks must be performed. LC/MS metabolomics data is inherently complex due to the presence of contaminants, adducts, isotopes, and other sources of chemical noise that do not provide unique biological information. These sources of noise must be removed to avoid erroneous results. Contaminants introduced during sample preparation or LC/MS analysis are annotated through comparison of the intensity of detected features between the method blank and pooled QC samples. Only features showing an intensity at least 3x greater in the QC than in the blank are retained for downstream analysis, as those signals with near equal intensity in the blank and QC samples are likely contaminants.^24^ To annotate degeneracy, MassID utilizes the mz.unity algorithm,^33^ which performs a combinatorial search of the *m/z* values of features that elute closely together and have correlated peak profiles. It also looks for known mass differences from possible isotopes, neutral losses, and oligomerization events with other metabolites and annotates these relationships to form a graph, which is to select a single preferred feature (monoisotopic protonated or deprotonated ion) amongst all detected signals of a metabolite to use as the reference signal. Previous work has shown that blank subtraction and mz.unity achieve noise removal comparable to experimental techniques, such as credentialing.^24^ However, as these tools are computational, they enable accurate noise filtration of any sample, not only those samples derived from specific experimental conditions (e.g., cells grown on isotopically labeled carbon sources).

### Metabolite Identification

The peaks surviving noise filtering are then subjected to metabolite identification using DecoID2, a novel, probabilistic algorithm that assigns a posterior probability of a metabolite identification being correct to each database match. DecoID2 is the successor to our previously reported DecoID software that performs a database-assisted deconvolution of MS/MS spectra.^34^ DecoID2 retains the MS/MS deconvolution functionality but leverages a novel scoring approach to metabolite identifications that generalizes beyond MS/MS data. This approach was inspired by a recent perspective^35^ that introduced the concept of identification probability. We have formalized a methodology to compute these probabilities and assembled a comprehensive metabolite-centric database of over 280,000 compounds complete with fragmentation data and retention times for >99% of all entries. The output of DecoID2 is a list of metabolite identifications and associated probabilities for each detected metabolite. Further description of the MassID database and DecoID2 are included in the subsequent sections.

### Metabolite Aggregation and Filtering

Lastly, all compounds showing a coefficient of variation (CV) greater than 25% are filtered to remove analytes with poor reproducibility. Compounds identified with DecoID2 can also be filtered to a defined false discovery rate (FDR). Unidentified metabolites are also filtered based on in-source fragment annotation (see Methods), a more stringent 10% CV filter, and removing metabolites whose *m/z* values do not match any compound in our metabolite database, as these likely reflect unannotated sources of noise or informatic artifacts. Identified metabolites and unknown compounds surviving filtering are then combined to form the metabolic profile of each sample.

### MassID Database

To identify metabolites, MassID leverages a comprehensive database of ∼280,000 compounds, complete with MS/MS fragmentation data for >99% of compounds. The data is from both proprietary in-house data as well as public sources, including HMDB^14^, Lipid Maps^36^, RefMet^37^, the Global Natural Products Social Molecular Networking database (GNPS)^38^, as well as prediction engines such as CFM-ID 4.0^39^ and NPClassifer^40^. The database includes many pieces of metabolite metadata and identifiers (e.g., SMILES, InChIKey, HMDB ID, PubChem CID, compound class/superclass/subclass, etc.) to facilitate data harmonization as well as multiomic integration. In total, this database contains more than 1.8 million MS/MS spectra. In addition to the public MS/MS spectra, we have acquired MS/MS data in our laboratory for over 1,000 compounds across multiple instruments. Along with this internal MS/MS data, experimental retention times for these compounds across our RPLC and HILIC methods were recorded. These reference retention times are used with DecoID2 to build a retention time prediction model to enable accurate predicted retention times (<1 min) for all database compounds (Figure S1) for both RPLC and HILIC methods.

### DecoID2: Probability-Based Metabolite Identification

DecoID2 is a novel module within MassID designed to structurally identify a detected metabolite through comparison of experimental LC/MS/MS data to reference data in the MassID database. Importantly, the novel aspect of DecoID2 is that database hits are scored quantitatively to give a posterior probability that a given metabolite identification is correct. By returning probabilities as opposed to qualitative scoring levels, we enable greater data transparency and allow false discovery rate (FDR)-controlled filtering of metabolite matches, which is the standard in other ‘omics (such as proteomics and genomics) as well as weighted incorporation of metabolites into downstream analyses like pathway analysis that account for the uncertainty in each metabolite identification.

The workflow for DecoID2 is depicted in Figure 3a. DecoID2 takes as input LC/MS/MS data as well as peak lists for the detected features for the study. First, the MS/MS and MS1 spectra for each feature are extracted. The MS/MS data is deconvolved with a database-assisted LASSO regression, as is also performed in DecoID^34^. After deconvolution, the *m/z* value of the feature is used to query the MassID database to gather candidate metabolite identifications. The deconvolved MS/MS spectra along with the MS1 spectrum for the query feature is then compared against the reference data in the MassID database (reference MS/MS data and theoretical isotope patterns) and the spectral similarity is scored via entropy similarity^41^. To enable usage of RT (retention time) for metabolite identification, our in-house library that includes RT data is also provided to the DecoID2 software. Based on the reference RTs, a random forest model is trained that predicts reference RTs for each assay individually based on the structure of the compound.^42,43^ The RT deviation for each potential metabolite identification is then tabulated. As a baseline, each set of potential metabolite identities for a given feature also includes an “unknown”, which accounts for the possibility that the metabolite’s true identity is not included in the MassID database. This “unknown” possibility is assigned the median similarity score across all features and potential identifications for each similarity type individually (MS1, MS/MS, and RT).

**Figure 3.**
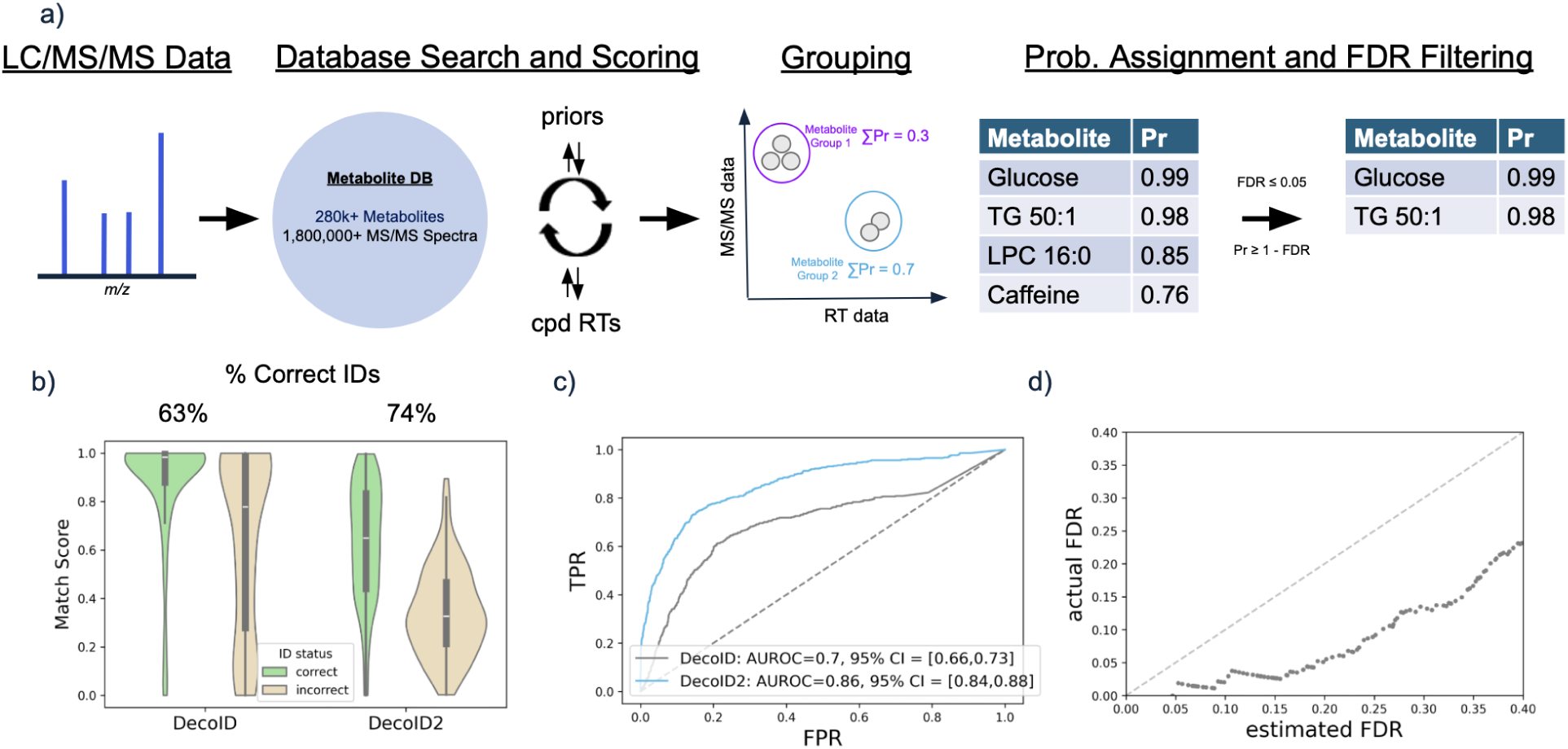
DecoID2 workflow and validation. a) LC/MS/MS are first searched against the MassID database. The acquired MS/MS are deconvolved prior to database matching. Predicted RT information is also used to score metabolite hits. Similarity scores (MS1, MS/MS, RT) are then combined in a Bayesian model to calculate metabolite identification probabilities. Then, metabolites are grouped based on if the similarity of reference data for potential metabolite IDs meet specified thresholds. Metabolite groups are then assigned a composite probability suitable for downstream analysis. b) Matching scores for the top hits when using MS/MS matching scores (DecoID) or identification probability (DecoID2) are shown in the violin plots when querying a LC/MS/MS dataset of a mixture of chemical standards. The percentage of top hits that correspond to correct identifications is listed above the DecoID and DecoID2 distributions. DecoID matching scores are high for both correct and incorrect hits. DecoID2 probabilities show greater separation. c) ROC curves for DecoID and DecoID2 scoring for the standard mixture dataset. d) DecoID2 estimated FDR versus actual FDR across different score cutoffs. DecoID2 estimated FDR is well correlated with the actual FDR and systematically overestimates the FDR.

The probability metric that underpins DecoID2 is used to compute the posterior probability that a metabolite identification, *m*, is correct given the degree of match to the available experimental data (*θ*) relative to all other potential metabolites in our database (*M*) as well as prior metabolite identification probabilities for all compounds (*γ*). This will be accomplished through Bayesian modeling of metabolite identification probabilities (Eq. 1).

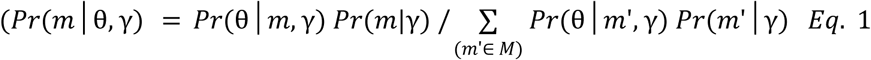

Here, θ corresponds to the degree of MS/MS match (α), degree of retention time match (β), and degree of natural abundance isotope pattern match (ζ). As these metrics come from independent sources and are independent of prior identifications, we decompose *Pr(θ│m,γ)* as *Pr(α│m) * Pr(β│m) * Pr(ζ│m)*. We compute *Pr(α│γ)*, *Pr(β│γ)*, and *Pr(ζ│γ)* after computing similarity metrics for each piece of experimental data. Intuitively, *Pr(α│γ), Pr(β│γ),* and *Pr(ζ│γ)* represent the probabilities of achieving these similarity scores given the metabolite identification is correct. Accordingly, we construct probability density functions (PDFs) for each type of similarity data by comparing similarity scores for metabolite identifications confirmed with orthogonal data (e.g., in-house metabolite library RT matches for building MS/MS PDF) or where a single potential metabolite identification was returned for a feature. These PDFs enable conversion of similarity scores into *Pr(α│γ), Pr(β│γ), and Pr(ζ│γ)* values. The prior metabolite identification probabilities are initialized based on confirmed metabolite identifications from diverse in-house data and updated based on maximum likelihood estimation periodically. In total, this approach assesses the degree of matching for each feature against all possible metabolite identities, integrates these matching data into a likelihood score of each potential identification for the feature, and then normalizes these scores by the sum of all likelihood scores for the potential identities. This ensures that the sum of identification probabilities for a query metabolite equals 1.0.

The last step in DecoID2 is what we refer to as metabolite grouping. Metabolite grouping takes inspiration from proteomics where proteins that cannot be differentiated based on their observed peptides are grouped together into what is referred to as a protein group.^44^ In metabolomics, we have a similar challenge in that metabolites may not be resolvable by MS1, MS/MS, or RT and would therefore show up as a single peak in the data. Traditionally, metabolite identifications are reported for the top match for the metabolite and ignore the possibility of multiple unique compounds being measured as a single signal. In addition, if two metabolites share the same reference data, they will achieve near identical matching scores, thus the selection of the best metabolite identification becomes arbitrary and can lead to erroneous results.

For DecoID2, these unresolvable compound identifications reduce the final identification probability, as DecoID2 probabilities are normalized to the total number of matches in the database. While correct, this becomes problematic for some types of compounds where many unresolvable compounds exist. One prominent example of this is complex lipids (e.g., triglycerides, diacylphosphodidylcholines, etc.) where you have a head group attached to one or more acylchains, where the acylchains can vary in terms of the number of carbon atoms and the number/position of double bonds. Without employing specialized methods, the exact position of double bonds within acyl chains of complex lipids cannot be resolved with LC/MS/MS. As many different forms of a lipid with a specified head group and acyl chain length exist, returned DecoID2 probabilities are very low as these lipid isomers will have near identical or identical reference data in the MassID database.

To address this issue, DecoID2 implements metabolite grouping. In this step, the potential identities of a metabolite are reviewed and metabolites sharing nearly identical reference data (same molecular formula, MS/MS, and predicted RT) are grouped together and their combined probability is summed. To facilitate a clear nomenclature, each metabolite in the MassID database contains a simplified name. This name is a more generic form of each metabolite that will be shared across multiple compounds and is sourced from lipid maps abbreviation^36^ and the RefMet reference names.^37^ For example, the triglyceride TG(16:0/16:0/18:1(11E)), containing two 16:0 acyl chains and an 18:1 (1 double bond) acyl chain, has the simplified name TG 50:1 (50 carbons and 1 double bond across all acyl chains). In many cases, the metabolite grouped together will have the same simplified name, allowing for that metabolite group to be clearly labeled. DecoID2 returns this grouped metabolite output along with the individual metabolite-level probabilities to facilitate downstream applications.

Prior to outputting the metabolite identifications, DecoID2 can also perform FDR filtering of hits if desired. For a single metabolite identification, the estimated FDR is simply 1 minus the identification probability. Therefore, for any given FDR target, by only taking metabolite identifications with probability greater than 1 minus the FDR target we can control the global metabolite identification FDR and ensure that the global FDR is less than or equal to the FDR target. In the example provided in Figure 3a, with a 5% FDR target, we will remove identifications having a probability <0.95, leaving two of the four compounds: glucose and TG 50:1.

### MassID Validation

To validate the MassID pipeline and DecoID2, we first leveraged a mixture of 455 detectable authentic metabolite standards across diverse chemical classes to mimic a complex biological sample while maintaining a known ground truth metabolome. We validated the detection and identity of these compounds when analyzed with our standard analytical LC/MS methods through manual data curation. After analysis, the resulting data (including MS/MS data) was processed with MassID. In total, 3,506 and 5,314 peaks were detected in the negative-mode and positive-mode dataset, respectively. After filtering for contaminants and redundancies, this was reduced to 337 and 250 peaks. When considering peaks corresponding to the standards, 355 of the 455 standards were successfully detected by MassID during untargeted peak detection and successfully survived noise filtering, demonstrating a sensitivity of 78.4%. For the other 8,352 peaks not corresponding to the reference signal of the chemical standards, 8,233 were successfully filtered, demonstrating a specificity of 98.6%. Software parameters can be tuned to increase sensitivity or specificity. For routine usage, however, we leverage parameter sets to maximize sensitivity to reduce the likelihood of false positives and erroneous results.

Next, we leveraged the standard dataset to evaluate the metabolite identification efficacy of DecoID2 relative to standard MS/MS matching after deconvolution, as is performed in the original DecoID software. After searching and scoring the filtered features against the MassID database, we evaluated the effectiveness of DecoID2’s probability metric for improving metabolite identification accuracy (the fraction of correct metabolite identifications returned as the top hit). Here, we found that DecoID2 achieves 74% accuracy across all compounds in the standard mixture, improving upon 63% accuracy with DecoID. Figure 3b shows the match scores for these incorrect and correct top hits for both DecoID (entropy similarity) and DecoID2 (identification probability) in a violin plot. As is apparent from the distribution, beyond the improvement in accuracy, a primary benefit of DecoID2 is increasing the separation of matching scores for correct and incorrect hits. With standard MS/MS matching in DecoID, both correct and incorrect identifications achieve a median entropy similarity of greater than 0.85, which is considered a strong match by most investigators.

To quantify the impact of the DecoID2 probability score on providing better separation between correct and incorrect metabolite identifications, we performed receiver operating characteristics (ROC) curve analysis on all hits for each of the metabolite standards detected. The ROC curve plots the true positive rate (TPR) versus the false positive rate (FPR) for all possible score thresholds for a given metric. An area under the ROC curve (AUROC) of 1.0 indicates perfect separation between incorrect and correct identifications while an AUROC of 0.5 indicates random performance. By comparing the distribution of scores for correct and incorrect identifications and computing the AUROC based on the returned DecoID2 probabilities and DecoID MS/MS similarity, we found that DecoID2 provides an AUROC of 0.85 (95% CI = [0.83, 0.87]). This is a significant increase over the AUROC of 0.70 (95% CI = [0.67, 0.73]) achieved by DecoID, demonstrating improved discrimination between correct and incorrect identifications with DecoID2 (Figure 3c).

The overall goal of DecoID2 is to enable quantitative assessment of metabolite identification confidence (the overall probability the metabolite identification is correct). This would enable FDR-controlled filtering of metabolite hits as well as incorporation of identification uncertainty into downstream analyses, instead of strict, somewhat arbitrary, filtering of hits based on qualitative confidence levels. To evaluate the ability of DecoID2 to perform FDR-controlled filtering, we resampled and applied different probability thresholds to the standard mixture dataset. We then calculated the estimated FDR for each dataset by subtracting the summed probability across all metabolites from the total number of metabolites. In addition, as the true identities of each metabolite are known, we can calculate the actual FDR by counting the number of incorrect identifications and dividing that by the total number of metabolites. We find that DecoID2-estimated FDRs are well correlated with the actual FDR but DecoID2 systematically overestimates the FDR (Figure 3d). While ideally DecoID2 FDR estimates would be exact against the actual FDR, an overestimate is preferred as it is a more conservative estimate.

## Application to Human Plasma

### Feature Detection and Metabolite Annotation

To evaluate MassID on a biological dataset, we next applied our metabolomics platform to human plasma samples from healthy individuals and individuals with cardiovascular disease (CVD). In total, 195,593 features were detected across all four assays. The breakdown of these features is provided in Figure 4a. Through data annotation for contaminants, artifacts, and common degeneracies, 97.14% of these features were annotated as noise, and 4,504 metabolites were structurally identified with an assigned identification probability from DecoID2. Of these, 4,090 metabolites were found from the untargeted analysis while the remaining 414 came from the targeted analysis. In addition, 1,366 features were annotated as in-source fragments. While these in-source fragment features are a relatively small portion of the total signals detected, they represent nearly 25% of the features remaining after standard data annotation/filtering. Without removal, these compounds would be assumed to be potentially novel metabolites and inflate the number of unknowns reported. Removal of these in-source fragments reduced the number of unidentified metabolites from 1,499 to just 133 compounds. In total, this amounts to near complete data annotation with 99.93% of features being either structurally identified or annotated as a defined noise signal.

**Figure 4:**
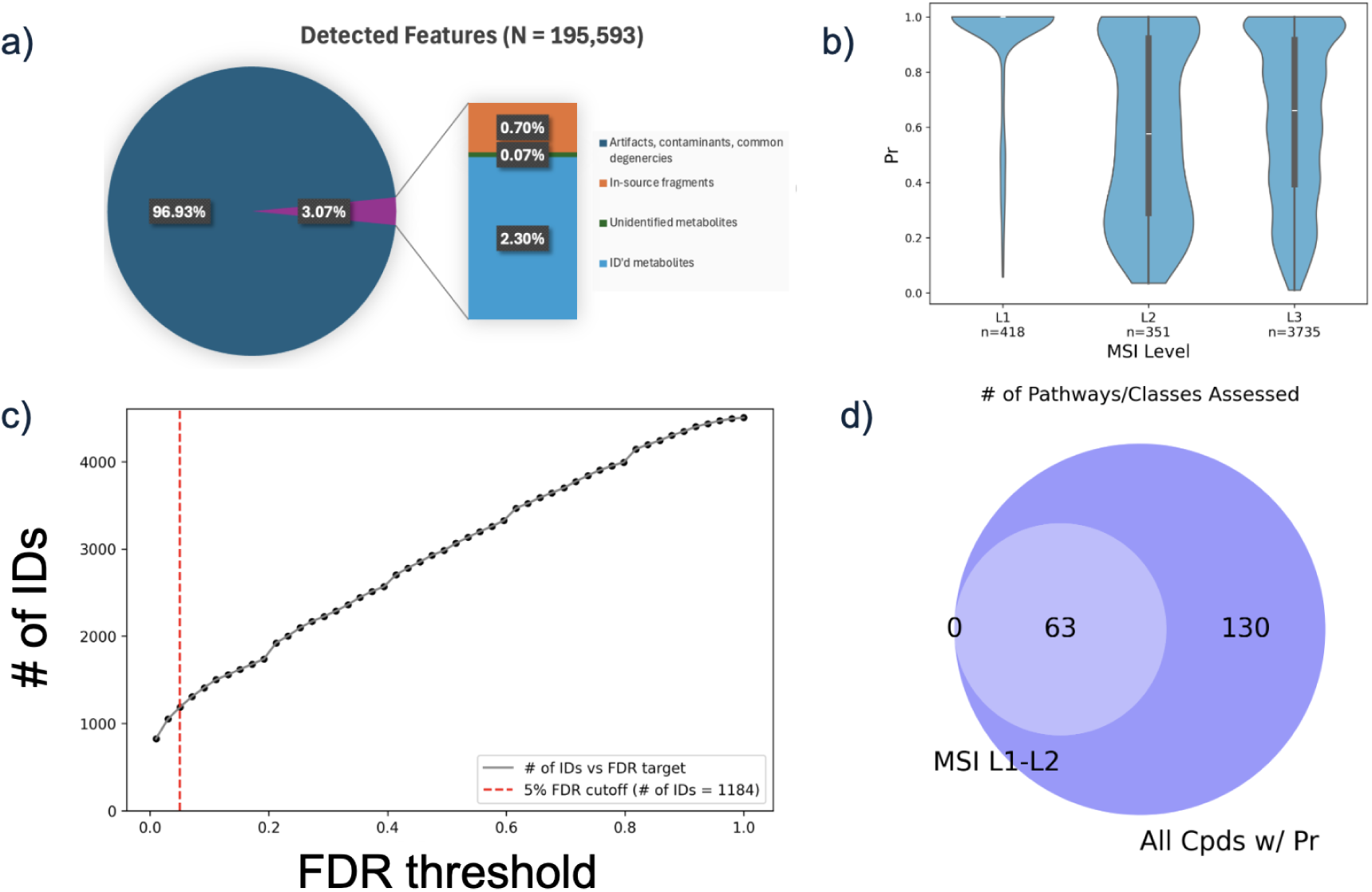
Application of MassID to human plasma. a) Breakdown of detected features in human plasma samples. b) Violin plots of metabolite identification probability versus MSI level shows moderate correlation with MSI levels. c) Scatter plot showing the number of identifications made at various FDR targets. d) Venn diagram showing the overlap of pathways that can be assessed when considering all identified metabolites (blue) or just those covered by MSI L1-L2 metabolites (grey).

### Comparing Identification Probabilities at the MSI Levels

Next, we sought to compare DecoID2 identification probabilities with the traditional MSI level classification scheme (see Methods). Traditionally, MSI level 1 and level 2 compounds are those used for downstream biological interpretation of results. In this human plasma dataset, 767 of the identified compounds met the criteria for such an identification and would thus be used to interpret results, discarding information from over 3,700 compounds. Figure 4b shows the distribution of identification probabilities for each MSI level. As expected, MSI level 1 and level 2 compounds (those with the most experimental evidence to support the identification) had higher overall probabilities. However, within each MSI level there is a dramatic range of identification probabilities. Notably, only 356/418 of MSI level 1 compounds were identified with >95% confidence, leaving 884 MSI level 2 and level 3 compounds that were identified at >95% confidence. This is predominately due to MSI levels not accounting for other potential metabolite identifications that may share high matching scores, while DecoID2 models the number of alternative matches along with the matching scores. In total, of the 4,504 metabolites profiled, 1,241 metabolites were identified with >95% confidence and the median identification probability across all identified metabolites was 69.3%, Figure 4c.

### Integrating Identification Probabilities into Pathway Analysis

To aggregate information from untargeted metabolomics analysis, pathway analysis is often performed. Traditionally, pathway analysis in metabolomics is limited to only highly confident (MSI L1-L2) metabolites, each given equal weight.^45^ This limits metabolome coverage as weakly identified metabolites are excluded. Other approaches have attempted to expand coverage by mapping all possible metabolite identifications onto pathways simultaneously.^46^ While this does expand pathway coverage, it introduces considerable noise as identifications are not weighted and metabolites have dramatic differences in the number of potential identifications possible. To address these limitations, we created a modified GSEA-based^47^ algorithm that accounts for identification confidence. Details of the methodology are provided in the methods. In brief, metabolite fold-changes, p-values, and identification confidence are used to sort metabolites into a ranked list, where the most upregulated, high confidence identified metabolites are at the top of the list and the most downregulated, high confidence identified metabolites are at the bottom. The algorithm assesses if metabolites in the pathway are clustered near the top or bottom of the ranked list. When applied to the plasma CVD dataset, this enabled 193 Reactome^48^ pathways and Lipid Maps^36^ classes were assessable (3+ metabolites mapped). As a comparison, we also mapped just the MSI L1 and L2 compounds onto the same pathways, which resulted in only 63 pathways that could be assessed, highlighting the dramatic increase in discovery potential made possible by MassID (Figure 4d).

### Assessing Metabolic Perturbations in CVD

Beyond these metrics, we also leveraged the metabolic profiles returned by MassID to uncover metabolic perturbations in CVD. To ensure reproducible results, we first inspected the technical variation present in the data through the calculation of CV values for all metabolites based on the calculated metabolite intensities in each of the QC samples used in this study. The median CV across the QC samples analyzed in this study was 3.4%. Conversely, when examining the research samples and tabulating the biological CV, we see a median biological CV of 47.1%, suggesting strong metabolic variation between individuals, Figure 5a.

**Figure 5:**
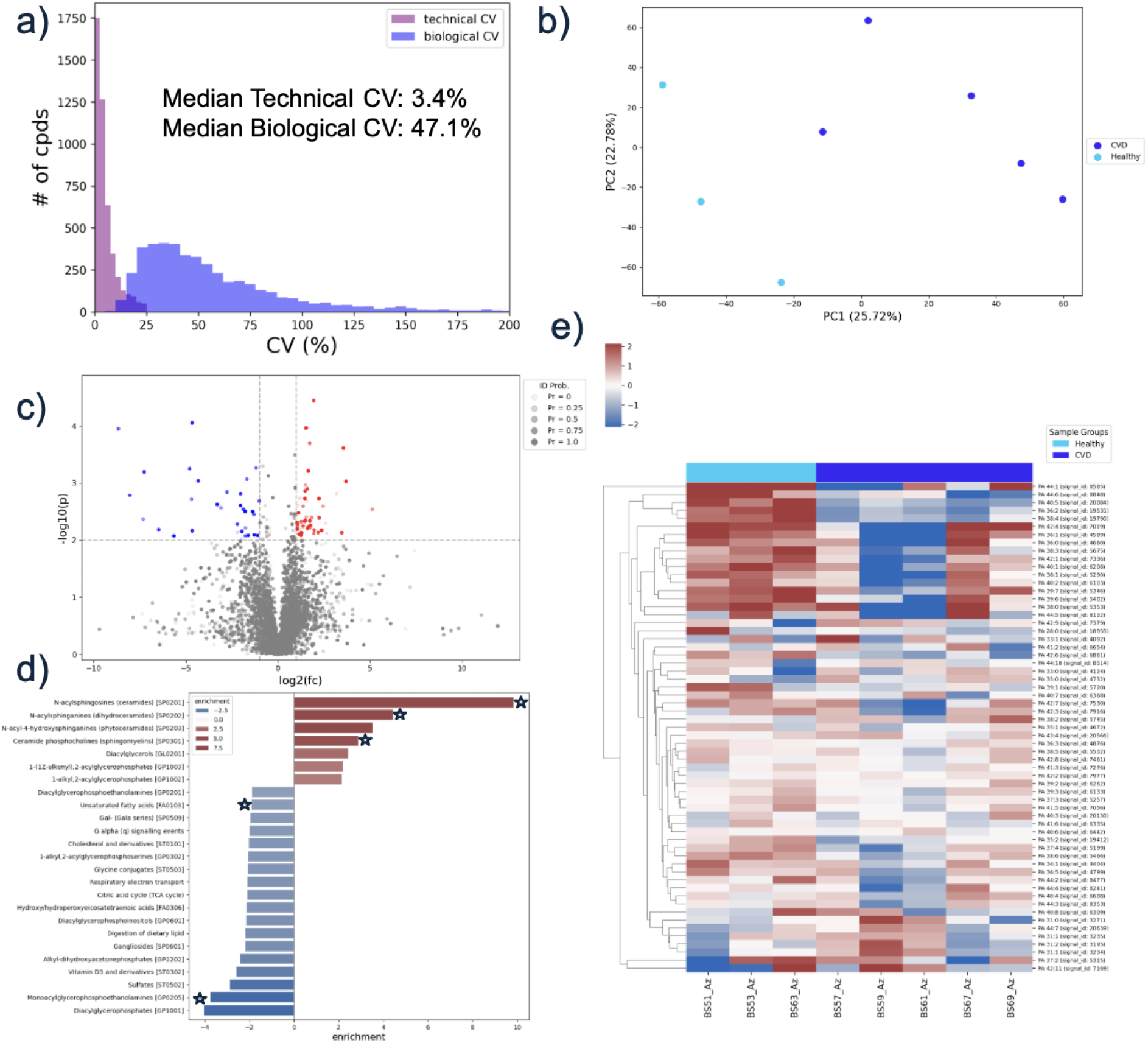
Metabolic Perturbation in CVD. a) Comparison of biological and technical CV. The histograms show the CV distribution from analysis of QC samples (purple) and research samples (blue). b) PCA plot of metabolic profiles, each dot is a sample. Dots are colored according to the experimental group. c) Volcano plot from statistical analysis showing log2(fc) and-log10(p) values. Each dot is a metabolite. Dots are shaded according to the identification probability. d) Result of pathway analysis. Each bar is a pathway or lipid class. The x-axis and bar color indicates the enrichment. Positive (red) values indicate metabolites in the pathway/class were overall increased in CVD relative to healthy samples. Negative (blue) values indicate overall decreased levels of the metabolites in the pathway/class. Starred pathways also reached statistical significance in the MSI L1-L2 analysis. g) Heatmap of PA lipids across sample groups. Cell color indicates the log2(fc) relative to the mean metabolite level across all samples.

Next, to capture global differences in the metabolic profiles between samples, we performed unsupervised principal component analysis (PCA, Figure 5b). Here we see that plasma samples generally cluster according to CVD and healthy groups, further indicating broad metabolic differences between the sample groups. To understand the specific metabolites driving this separation, we leveraged one-way Welch’s analysis of variance (ANOVA) to identify metabolites showing a statistically significant difference in abundance between healthy and CVD individuals. The analysis highlighted 24 metabolites identified at >95% confidence showing a statistically significant difference in metabolite abundance (p<0.05, |log2(fc)|>1), Figure 5c. The most differentially abundant metabolite was the very long chain polyunsaturated fatty acid, FA 38:4, that was increased in CVD patients, which aligns with the known dysregulation of lipid metabolism in CVD. Elevated levels of certain fatty acids can contribute to lipotoxicity, inflammation, and oxidative stress, all of which are implicated in the progression of CVD.^49,50^

To assess metabolic differences at the pathway level, we performed the weighted GSEA-based algorithm described in the section above, resulting in 25/193 pathways and lipid classes assessed showing significant differential enrichment (Figure 5d). Notably, when considering only MSI L1 and L2 compounds, only 5 pathways showed differential enrichment. It is important to note that all 5 pathways were also differentially enriched when leveraging all identified compounds, weighted by their probability, serving as validation of the approach.

The most enriched pathway/class was ceramides, which was also 1 of the 5 pathways found to be significantly enriched when only considering MSI L1-L2 compounds. Ceramides are a class of lipids that play crucial roles as structural components of cell membranes and as signaling molecules. They are involved in various cellular processes, including inflammation, insulin resistance, endothelial dysfunction, and programmed cell death (apoptosis). Dysregulation of ceramide levels has been observed in various disease states, including cardiovascular disease. The increase in ceramide levels was quite uniform across the class and consistent within CVD and healthy groups.

The most decreased pathway/class was diaglycerophosphates (i.e., PAs), Figure 5e. Notably, this class was missed in the analysis considering only MSI L1-L2 compounds, highlighting a missed insight that can be recovered by leveraging identification probabilities. The chronic inflammation and oxidative stress characteristic of CVD can disrupt the normal activity of enzymes that produce PA lipids, such as phospholipase D.^51^ Furthermore, the increased metabolic demands and cellular stress associated with heart disease can lead to the rapid conversion of PA into other signaling lipids or its degradation to fuel other cellular processes, thereby lowering its overall concentration. These changes in PA levels can, in turn, affect critical cellular functions, including signaling pathways that regulate cell growth, proliferation, and survival, potentially contributing to the progression of cardiovascular disease.

Many other lipid classes were also elevated, including other sphingolipids and diaglycerols. In terms of metabolic pathways, of note were the decreased levels of many TCA cycle intermediates, such as fumarate and malate, that drove the statistical negative enrichment of the TCA cycle. TCA cycle activity is dependent on mitochondrial activity, which is known to be impaired in CVD.^52^

## Conclusion

Untargeted metabolomics offers unparalleled insights into cellular processes and disease states. However, the complexity of untargeted metabolomics data, characterized by pervasive noise and challenges in accurate metabolite identification, has hindered its broad application in biomedical research. To address these limitations, we developed a workflow that leverages four complementary LC/MS assays (RPLC and HILIC in both positive and negative ionization modes) and an analysis pipeline, MassID, that enables comprehensive metabolome analysis.

The MassID software integrates a suite of sophisticated software functionalities, including deep learning-based peak detection (PeakDetective), comprehensive noise filtering (mz.unity and contaminant removal), and probabilistic metabolite identification (DecoID2). These components work in concert to address the intrinsic challenges of untargeted metabolomics. Unlike traditional qualitative scoring systems, DecoID2 assigns a posterior probability to each metabolite identification, providing a quantitative assessment of its correctness. This quantitative probability enables FDR-controlled filtering of metabolite hits and allows for weighted incorporation of metabolites into downstream analyses, akin to best practices in proteomics and genomics.

Furthermore, DecoID2 addresses the challenge of unresolvable metabolite isomers (e.g., complex lipids) by implementing a metabolite grouping strategy, which combines compounds sharing identical reference data and sums their probabilities, providing a more chemically relevant output. Comparison of DecoID2 probabilities with traditional MSI levels shows moderate correlation between identification confidence and MSI level. However, wide probability distributions occur within each MSI level, demonstrating the importance of quantitative scoring that factors in both matching scores and alternative hits. Only 85% of MSI level 1 compounds were identified at <5% FDR and the majority of <5% identifications came from MSI level 2 and level 3 compounds.

Applying MassID to human plasma samples from healthy and CVD individuals showcased its ability to achieve near-complete data annotation. MassID successfully filtered out ∼97% as contaminants, degeneracies, or artifacts, and an additional 0.7% as in-source fragments. This resulted in the measurement and structural identification of over 4,500 unique metabolites with quantified DecoID2 probabilities, leaving less than 0.1% of features unannotated. Of these >4,500 compounds, 1,241 were identified with >95% confidence. Comparative analysis of the metabolic profiles of CVD and healthy individuals show key metabolic changes in specific compounds as well as lipid classes and metabolic pathways, such as ceramides, PA lipids, and the TCA cycle, recapitulating many known metabolic consequences of CVD.

In conclusion, MassID represents a significant leap forward for untargeted metabolomics. By addressing the challenges of noise, identification, confidence, and scalability, MassID provides a truly robust, accurate, and cloud-based solution for biochemical discovery. This platform is poised to power genuine small molecule discovery, facilitate novel biological insights, and accelerate advancements in life-science research.

## Methods

### Standard Mixture Validation Dataset

Data from authentic standard mixture were generated from samples prepared and analyzed as described in the original DecoID publication. Metabolite standards were sourced from IROA technologies. Standards were analyzed via HILIC/MS/MS as described below.^34^

## Sample Preparation

### CVD Plasma Samples

A 50 µL aliquot of plasma was transferred onto a solid phase extraction (SPE) system and 10 µL was taken from each sample to form the pooled quality control sample. 200 µL of 1:1 acetonitrile:methanol (ACN:MeOH) was added to each well and shaken for 1 min at room temperature at 360 rpm, followed by a 10 min incubation. Next, 150 µL of 2:2:1 MeOH:ACN:H_2_O was added to each well and taken again for 10 min. Polar metabolites were then eluted into the 96-well collection plate by using the positive pressure manifold. This is repeated with 100 µL of 2:2:1 MeOH:ACN:H_2_O into the same collection plate. Polar eluates were then covered and stored at-80°C until LC/MS analysis.

The SPE plates from the polar metabolite extraction were washed twice with 500 µL 1:1 methyl tert-butyl ether:MeOH to elute lipids into a new collection plate using a positive pressure manifold. The combined eluates were dried under a stream of nitrogen at room temperature and reconstituted with 200 µL 1:1 isopropanol (IPA):MeOH prior to LC/MS analysis.

## LC/MS Data Collection

### LC/MS analysis of polar metabolites

LC/MS mobile phases A and B were prepared as follows: A) 20mM ammonium bicarbonate, 0.1% ammonium hydroxide, 5% ACN, 2.5 mM medronic acid and B) 95% ACN.

A 2 µL aliquot of polar metabolite extract was analyzed with HILIC/MS by using the following linear gradient at a flow rate of 250 µL/min: 0-1 min: 90% B, 1-12 min: 90-35% B, 12-12.5 min: 35-20% B, 12.5min-14.5 min: 20% B. The column was re-equilibrated with 20 column volumes of 90% B. Mass spectrometry analysis was completed with a mass range of 67-1500 Da with 1 scan/sec and a mass resolution of 120,000 in both positive and negative ionization mode. MS/MS data were acquired in a data-dependent iterative fashion with a 1.3 m/z isolation window.

### LC/MS analysis of lipid metabolites

LC/MS mobile phases A and B were prepared as follows: A) 5:3:2 H_2_O:ACN:IPA, 10 mM ammonium formate, and 5 μM Agilent deactivator additive and B) 1:9:90 H_2_O:ACN:IPA, 10 mM ammonium formate.

A 4 μL aliquot of lipid metabolite extract was analyzed with RPLC/MS (Waters Acquity Premier HSS T3, 2.1 x 100 mm) by using the following linear gradient at a flow rate of 400 μL/min: 0-2.5 min: 15-50% B, 2.5-2.6 min: 50-57% B, 2.6-9 min: 57-70% B, 9.0-9.1 min: 70-93% B, 9.1-11 min: 93- 96% B, 11.0-11.1 min: 96-100% B, 11.1- 12 min: 100% B, 12.0-12.2 min: 100-15% B.

The column was re-equilibrated for 3.8 min. Mass spectrometry analysis was completed with a mass range of 100-1700 *m/z* with 1 scan/sec in both positive and negative mode. MS/MS data was acquired in a data-dependent iterative fashion with a 1.3 m/z isolation window.

### Quality Control Processes

Quality control (QC) samples are analyzed at the start of the analytical run to stabilize the instrument and periodically throughout the experiment to assess system stability and address sources of experimental variability during data processing.

### Sample Preparation

During metabolite extraction, stable isotopically labeled metabolites are added to the extraction solvent(s). For polar metabolite analysis, a mixture of 20 fully labeled (^13^C and ^15^N) amino acids are utilized. For lipid analysis, a mixture of 14 deuterated lipids is utilized that span 14 different lipid classes. The addition of these internal standards enables QC of metabolite extraction, recovery, as well as downstream data quality post LC/MS data acquisition.

During the initial sample processing, a small aliquot of each individual sample (prior to metabolite extraction) is reserved to combine for a reference pooled QC sample. The amount of pooled QC sample generated is proportional to the total number of analysis batches for a study. These pooled QC samples are then aliquoted and frozen at-80 °C. For each analysis batch, a pooled QC aliquot is thawed and prepared with the analysis batch to account for batch effects introduced during sample preparation.

For each batch of samples that are prepared for metabolomics analysis, a method blank is also produced which is pure water that undergoes all steps of sample preparation and downstream analysis. This sample enables annotation of potential small molecule contaminants introduced during sample preparation and/or analysis.

### Data Acquisition

Prior to beginning data acquisition for a study, the mass spectrometer is calibrated to ensure mass accuracy and signal transmission. Additionally, a mixture of authentic standards is analyzed to assess chromatographic stability and ensure retention time drift of <30s. At the beginning of each analysis batch, three injections of a pooled plasma extract are used to condition the LC column and system.

In addition, both the method blank and pooled QC samples are analyzed at the beginning and end of each analysis batch. The pooled QC sample is also run in between every 10 samples to monitor for batch and run-order effects.

### Data Quality Assessment

The total ion current (TIC) of each sample is compared against the pooled QC TIC and blank TIC. Samples showing a greater overall similarity to the blank are flagged as these likely indicate a failure of either data acquisition or sample preparation. In addition, extracted ion chromatograms for all internal standards are extracted, and samples showing outlier intensity values are flagged. Samples deemed to have poor data quality after inspection are flagged, and samples are automatically queued for re-analysis on the LC/MS system.

### In-source Fragment Annotation

After metabolite identification, there are metabolites that are identified as well as unknowns. One difficult to annotate form of noise in LC/MS-based metabolomics are in-source fragments that when not removed inflate the number of unknowns present in a dataset.^36,47^ These are ions that are generated from fragmentation of other metabolites and occur inside the mass spectrometer. To annotate such fragments we leverage the comprehensive MS/MS data in the MassID database to annotate in-source fragments that appear within the list of unknown, potentially novel metabolites returned by DecoID2. The workflow is shown in Figure S2. After application of DecoID2, the list of unidentified and identified metabolites are leveraged together for fragment annotation. First, a custom database of fragment *m/z* values is gathered by querying the MassID database for low-energy predicted fragmentation spectra for identified metabolites. The *m/z* values of the predicted fragments along with the retention times of their corresponding metabolites identified in the sample are used to form a custom fragment database. These *m/z* values are retention times are then compared to the *m/z* values and retention times of the unidentified metabolites. Those with a match are then annotated as a fragment and can be removed prior to downstream analysis.

### Data Reporting and Filtering

Metabolite identifications are classified based on the Metabolomics Standards Initiative (MSI) scoring scheme^21^ with values ranging from Level 1 to Level 4. Level 1 through Level 3 identifications are all supported by accurate mass matches. Level 1 identifications are additionally supported by experimental retention times and MS/MS spectra from authentic standards. Level 2 identifications are made based on MS/MS matching to in-house or external reference data Level 3 identifications are made based solely on accurate mass. Level 4 identifications represent unique LC/MS signals that cannot be identified by database matching.

Metabolite signals are discarded if the CV among quality control samples is greater than 25%. Unidentified metabolites are further filtered if they have CV values greater than 10%, to ensure confidence that signals are not information artifacts. Batch and run-order effects are corrected for using a random forest algorithm trained on the drift observed in the QC samples.^26^ Metabolite intensities are log2 transformed prior to statistical analysis. Metabolite identifications and intensities are manually reviewed for concordance and accuracy.

## Statistical Analyses

Hypothesis testing was executed utilizing a one-way ANOVA, which does not assume equal variances among the groups. Log2 transformed metabolite intensities were used for null hypothesis testing. Fold-changes were computed from non-log2 transformed intensities.

To perform pathway analysis, we formed a ranked list of metabolites where metabolites with the most positive fold-changes are at the top and the most negative are at the bottom. We then applied the classical gene-set enrichment analysis (GSEA) algorithm^47^ to calculate the enrichment of each pathway based on the ranked list. Metabolites in the ranklist were weighted according to the identification probability. The statistical significance of each pathway enrichment score was assessed through shuffling the association between analytes and pathways and recalculating enrichment values 1000 times to form an empirical null distribution of pathway enrichment values. For this analysis, Reactome^48^ pathways were used as the pathway database and Lipid Maps^36^ was used for lipid class assignment.

## Supplementary Information

**Figure S1:**
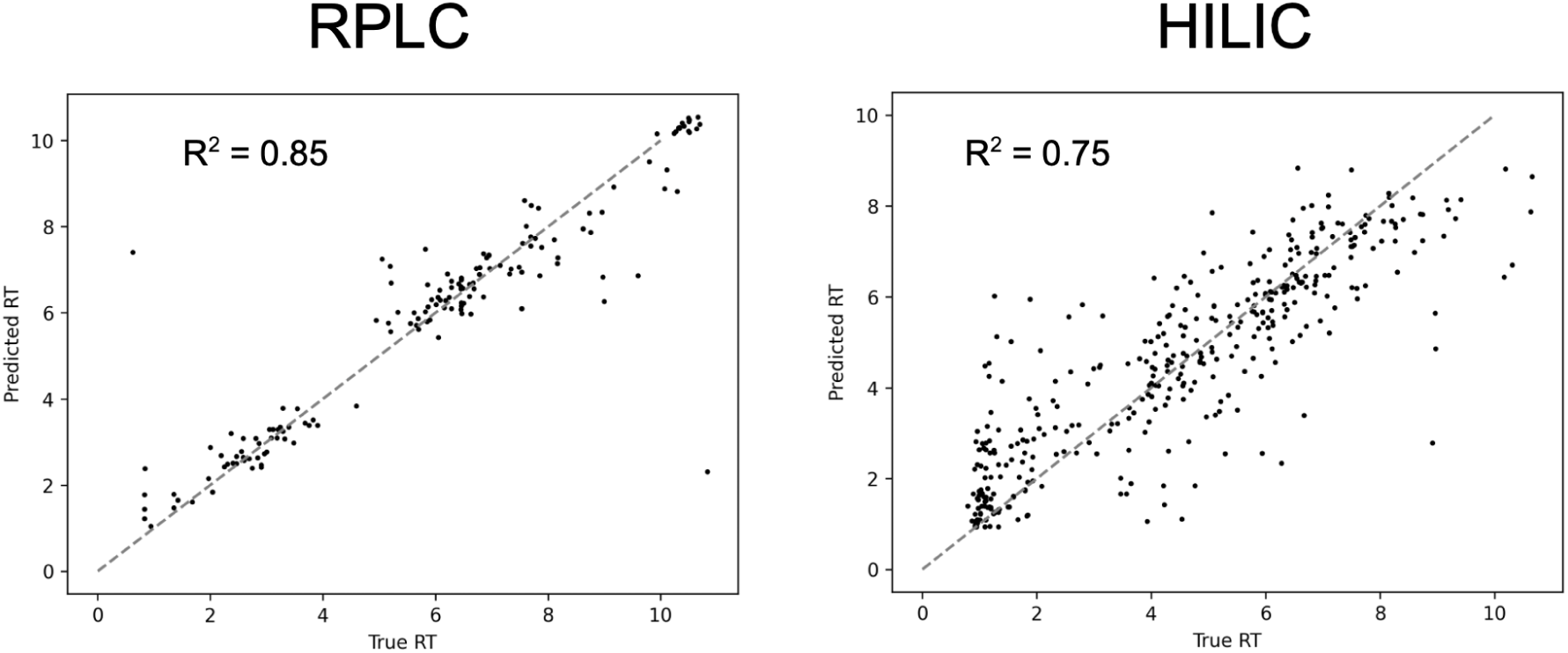
Validation of retention time predictions. The above scatter plots show the cross validated predictions of retention times for our RPLC method (left) and our HILIC method (right). The x-axis shows the true retention time (RT) of each compound in the library. The y-axis shows the predicted retention time from a RT model trained on held out data. Data are from 10-fold cross validation.

**Figure S2:**
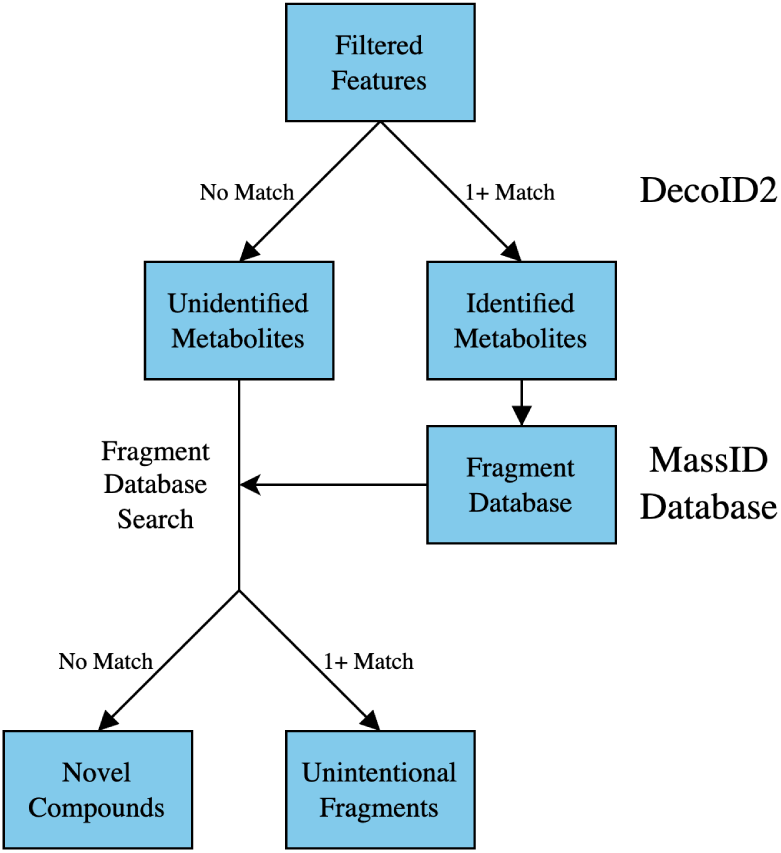
PeakDefrag workflow. After feature filtering, unidentified and identified metabolites are leveraged together to annotate signals in the remaining unknown metabolites that may be a fragment of another identified metabolite in the same experiment. The low-energy fragments of all identified metabolites are predicted, gathered, and compared to closely eluting and well correlated unidentified metabolites. Those with a *m/z* match are excluded from downstream analyses.

## Notes

### Competing Interest Statement

ES, AR, MG, AM, SA, DG, and TC are employees of Panome Bio Inc. GJP is the CSO of Panome Bio. KC is a consultant for Panome Bio. The Patti laboratory has a research collaboration agreement with Agilent Technologies.

